# Beclin-1 restrains aldosterone signaling via autophagic degradation of the mineralocorticoid receptor to protect against cardiovascular injury

**DOI:** 10.64898/2026.05.19.726128

**Authors:** Lei Wang, Wen-Yi Jiang, Hao-Tian Zhang, Xiao-Wei Sun, Ya-Mai Gao, Koji Murao, Guo-Xing Zhang

**Author notes:** Corresponding authors: Lei Wang Institute for Translational Neuroscience of Nantong First People’s Hospital, Affiliated Hospital of Southeast University, Nantong, 226014, P.R. China; Guo-Xing Zhang Department of Physiology and Neuroscience, Medical College of Soochow University, 199 Ren-Ai Road, Dushu Lake Campus, Suzhou Industrial Park, Suzhou, 215123, P.R. China. These authors contributed equally to the work (Lei Wang, Wen-Yi Jiang, Hao-Tian Zhang, Xiao-Wei Sun, Ya-Mai Gao).

## Abstract

Cells deploy adaptive programs to maintain homeostasis under stress, yet mechanisms counteracting damage triggered by transmembrane signaling remain poorly defined. Using a hyperaldosteronism model, we examined how autophagy regulates aldosterone-mediated mineralocorticoid receptor (MR) activation. In human umbilical vein endothelial cells (HUVECs), aldosterone induced autophagy, as evidenced by elevated Beclin-1, an increased LC3-II/LC3-I ratio, and reduced SQSTM1/p62. Aldosterone also promoted MR translocation from the cytosol to the nucleus. Co-immunoprecipitation and immunofluorescence revealed direct interaction and colocalization between MR and Beclin-1, as well as enhanced MR-lysosome association. Domain mapping showed that the Beclin-1 middle domain (161–241 AA) binds the MR C-terminal region (601–984 AA). Bioinformatic prediction and ChIP-qPCR confirmed that MR occupies the promoters of *IL-1β*, *IL-6*, and *TNF-α* upon aldosterone stimulation. Beclin-1 overexpression attenuated MR nuclear translocation, promoter binding, and inflammatory cytokine expression, whereas Beclin-1 knockdown reversed these effects. In vivo, aldosterone-infused Beclin-1 transgenic (*Becn1-tg*) mice exhibited lower blood pressure, reduced aortic medial thickening, and attenuated cardiac hypertrophy relative to wild-type controls, with no difference in body weight. Our findings identify Beclin-1 as a critical negative regulator of aldosterone signaling through an autophagy-dependent negative feedback loop. By interacting with MR and directing it toward lysosomal sequestration, Beclin-1 limits MR nuclear translocation and transcriptional activity, thereby mitigating aldosterone-induced vascular inflammation and cardiovascular injury.

**Highlights:**

Aldosterone activates autophagy and promotes MR–Beclin-1 interaction in HUVECs
Beclin-1 binds the C-terminal MR domain and directs MR to lysosomal degradation
Beclin-1 overexpression suppresses MR nuclear translocation and cytokine gene activation
Beclin-1 transgenic mice are protected from aldosterone-induced cardiovascular injury

## Introduction

Cells inhabit a dynamic and frequently hostile microenvironment, continuously challenged by diverse stimuli spanning soluble hormones, cytokines, mechanical stress and pathogenic factors. The plasma membrane serves as the primary interface, preserving cellular structural integrity and governing the regulated exchange of ions, metabolites and signaling molecules^1, 2^. Far from being a passive barrier, the membrane orchestrates environmental cue integration via an array of receptors and channels^3^. Transmembrane signaling is indispensable for cellular survival and adaptation, yet sustained or aberrant pathway activation disrupts homeostasis, elicits maladaptive stress responses and drives pathological progression^4, 5^. A central unresolved question in cell biology is how cells balance the benefits of environmental sensing with the risks of signaling overload.

Among endocrine signals governing cellular physiology, aldosterone has occupied a pivotal role for over six decades. Identified in the mid-20th century as a mineralocorticoid essential for sodium retention and potassium excretion, aldosterone is now recognized as a pleiotropic regulator with functions extending far beyond epithelial ion transport^6^. The canonical effector of aldosterone is the mineralocorticoid receptor (MR), a ligand-activated transcription factor belonging to the nuclear receptor superfamily. In its unliganded state, MR localizes to the cytoplasm in complex with chaperone proteins (including heat shock proteins and immunophilins) that stabilize its conformation^7^. Upon hormone binding, MR undergoes conformational rearrangement, dissociates from inhibitory complexes and translocates to the nucleus. There, MR binds to mineralocorticoid response elements (MREs) in target gene promoters, modulating transcriptional programs governing electrolyte transport, cell growth and stress responses^8^.

In addition to genomic actions, aldosterone rapidly modulates cellular activity via non-genomic signaling, potentially involving membrane-localized MR or crosstalk with MAPK, PI3K/Akt and PKC cascades^9, 10^. These rapid effects expand aldosterone signaling scope, enabling fine-tuned responses to acute physiological challenges, yet potentiate deleterious signaling under chronic hormone elevation.

Dysregulated aldosterone signaling is firmly established as a driver of cardiovascular pathology. Excess aldosterone (as in primary hyperaldosteronism) contributes not only to salt-sensitive hypertension but also to tissue remodeling, fibrosis, vascular stiffening and heart failure^11, 12^. Clinical and experimental evidence indicates aldosterone promotes oxidative stress, inflammation and extracellular matrix deposition in the cardiovascular system, accelerating target organ damage. In this context, cellular intrinsic stress-buffering capacity is overwhelmed, raising critical questions about failed cellular defense mechanisms under sustained MR activation^13^.

Autophagy, a highly conserved lysosomal degradation pathway, maintains cellular homeostasis by recycling damaged organelles, aggregated proteins and dysfunctional mitochondria^14^. As a core quality-control system, autophagy eliminates toxic components, particularly under metabolic or oxidative stress. Emerging studies reveal a complex, context-dependent relationship between aldosterone signaling and autophagic activity. In some settings, aldosterone-induced stress activates autophagy as a compensatory mechanism; in others, excessive or dysregulated MR activation impairs autophagic flux, exacerbating cellular injury^15^. This duality underscores autophagy as a critical node integrating adaptive and maladaptive signaling in cardiovascular tissues.

Elucidating crosstalk between aldosterone-MR signaling and autophagy provides a framework to explain cellular failure in defending against chronic hormone excess. MR-driven transcriptional programs may alter autophagy-related gene expression, while non-genomic cascades directly modulate autophagic flux via phosphorylation of key regulators (ULK1, Beclin-1, mTOR). Conversely, defective autophagy amplifies MR signaling by enabling damaged mitochondria accumulation and reactive oxygen species overproduction, creating a vicious cycle promoting tissue remodeling and dysfunction.

These insights carry pivotal translational implications. Hyperaldosteronism is increasingly recognized as a cause of resistant hypertension and an independent risk factor for arrhythmias, myocardial infarction and stroke^12, 16^. Current therapies (e.g., mineralocorticoid receptor antagonists) confer clinical benefit but are limited by side effects and incomplete efficacy^17^. Dissecting autophagy-mediated modulation of MR-driven pathology may uncover novel therapeutic targets, via pharmacologic activation of autophagic flux or combinatorial strategies attenuating MR signaling and downstream metabolic stress.

Collectively, cells maintain a delicate equilibrium between sensing and safeguarding, vividly exemplified by the dual nature of aldosterone signaling. While essential for fluid and electrolyte homeostasis, aldosterone drives cardiovascular disease in excess, in part by overwhelming cellular defense systems. Autophagy, a core homeostatic pathway, emerges as both a key mediator and therapeutic target in this process. Dissecting interplay between transmembrane signaling, MR activation and autophagic regulation yields fundamental insights into cellular adaptation under stress and opens new avenues for treating hyperaldosteronism-related cardiovascular disorders.

## Methods

### Protein Analysis in HUVECs

Human umbilical vein endothelial cells (HUVECs) were maintained in vascular endothelial cell growth medium supplemented with 10% fetal bovine serum (FBS, #11011-8615, Hangzhou Sijiqing Company), 100 U/mL penicillin (#C0222, Beyotime Biotechnology), and 100 µg/mL streptomycin (#ST487, Beyotime Biotechnology) at 37 °C in a humidified 5% CO₂ incubator. Upon reaching 80–90% confluence, cells were serum-starved for 12 h to synchronize metabolic activity before stimulation. Aldosterone (#A9477, Sigma) was then applied at concentrations ranging from 10^-9^ mol/L to 10^-5^ mol/L for durations of 1-24 h, as indicated.

Following treatment, cells were lysed on ice in radioimmunoprecipitation assay (RIPA, #P0013C, Beyotime Biotechnology) buffer supplemented with protease and phosphatase inhibitors. Total protein concentrations were quantified using the BCA assay (#P0010, Beyotime Biotechnology), and equal amounts of protein were resolved by SDS-polyacrylamide gel electrophoresis (SDS-PAGE) and transferred onto polyvinylidene difluoride (PVDF) membranes. Membranes were blocked in 5% non-fat milk in Tris-buffered saline containing 0.1% Tween-20 (TBST) for 1 h at room temperature, followed by overnight incubation at 4 °C with primary antibodies against Beclin-1 (#ab207612, Abcam), LC3-I/II (#ABC929, Sigma), and SQSTM1/p62 (#7695T, Cell Signaling Technology). After washing in TBST, membranes were incubated with horseradish peroxidase (HRP)-conjugated secondary antibodies (#GAM007, Multi Sciences) for 1 h at room temperature. Protein bands were visualized using enhanced chemiluminescence (ECL, #1705060, BIO-RAD) detection.

### Co-Immunoprecipitation (Co-IP)

HUVECs were transfected with plasmids encoding Myc-tagged MR and FLAG-tagged Beclin-1 using Lipofectamine 3000 (#L3000015, Thermo Fisher Scientific) according to the manufacturer’s protocol. After 48 h, cells were harvested and lysed in ice-cold immunoprecipitation (IP) buffer containing protease inhibitors. The lysates were precleared with protein A/G agarose beads (#16-663, Sigma) for 1 h at 4 °C and subsequently incubated with anti-FLAG antibodies overnight at 4 °C. Immune complexes were captured with protein A/G agarose beads for an additional 2 h at 4 °C, followed by extensive washing with IP buffer. Bound proteins were eluted by boiling the beads in SDS sample buffer and analyzed by western blotting using anti-Myc antibodies (#60003-2-lg, Proteintech).

### Plasmid Construction and Domain-Mapping Co-Immunoprecipitation

Full-length Beclin-1 cDNA was amplified from a human cDNA library and subcloned into a Myc-tagged vector. Full-length Beclin-1 cDNA was amplified from a human cDNA library and subcloned into a Myc-tagged vector. Full-length MR cDNA was cloned into a Strep-tagged vector. The truncated mutants were generated according to gene functional domains: Beclin-1 1-160 AA (112–159 AA: Bcl-2/Bcl-XL binding domain), 161-240 AA, and 241-450 AA (245–450 AA: evolutionarily conserved domain, 425–450 AA: membrane-association domain). MR 1-175 AA (1-169 AA: activation function 1a domain), 176-600 AA 451–602 (activation function 1b domain), 601-984 AA (603–669 AA: DNA-binding domain, 732–984 AA: activation function 2 domain). All constructs were verified by Sanger sequencing. Co-IP was performed using Myc magnetic beads (#B26301, Bimake), and bound proteins were detected by anti - Strep antibodies (#ab76949, Abcam). All constructs were verified by Sanger sequencing.

### Binding Site Prediction

Promoter sequences of *IL-1β*, *IL-6*, and *TNF-α* were retrieved from NCBI GenBank (https://www.ncbi.nlm.nih.gov). Putative MR-binding motifs were predicted using the JASPAR 2024 (https://jaspar.elixir.no/) database and validated by literature comparison.

### Genetic Manipulation and Functional Analysis of MR–Beclin-1 Signaling

To manipulate Beclin-1 expression, HUVECs were transduced with lentivirus carrying Beclin-1 cDNA for overexpression, or transfected with Beclin-1-specific siRNAs (sense: 5’-GCUCAGUAUCAGAGAGAAUTT-3’, antisense: 5’-AUUCUCUCUGAUACUGAGCTT-3’) using Lipofectamine 3000. At 48 h after transduction or transfection, cells were stimulated with aldosterone.

For immunofluorescence, cells were fixed, permeabilized, and incubated with anti-MR antibodies ((#ab64457, Abcam). Nuclei were counterstained with DAPI (#10236276001, Roche). Lysosomes were labeled with LysoTracker (#sc-18822, Santa Cruz).

ChIP-qPCR assays were performed to assess MR binding to *IL-1β*, *IL-6*, and *TNF-α* promoters.

### In Vivo Aldosterone Infusion and Physiological Assessment

Beclin-1 transgenic (*Becn1-tg*) mice and wild-type (WT) littermates were infused with aldosterone (280 µg/kg/day) or vehicle for 4 weeks using ALZET minipumps (#1004, Alzet). Systolic blood pressure was measured by tail-cuff. Aortas and hearts were harvested for histological and molecular analyses.

### Statistical Analysis

All data are expressed as mean ± S.E.M. Statistical comparisons between multiple groups were performed using ANOVA followed by Newman–Keuls post hoc tests, whereas comparisons between two groups were analyzed using unpaired two-tailed Student’s t-tests. P values < 0.05 were considered statistically significant. Statistical analyses and visualization were performed using R (version 4.3.2).

## Results

To develop an optimal experimental model in HUVEC, cells were treated with a series of concentration gradients of aldosterone across multiple time points. Aldosterone treatment (10^-7^ mol/L) elicited a striking temporal pattern of autophagy-related protein expression. Beclin-1 levels began to rise as early as 12 hours after exposure, showing a sustained and progressive increase thereafter (Ps < 0.05; Figure 1A). In parallel, the conversion of LC3-I to LC3-II became markedly elevated after 24 hours of treatment, indicating activation of autophagic flux (Ps < 0.05; Figure 1B). Concomitantly, the accumulation of SQSTM1/p62 declined significantly from 24 hours onward (Ps < 0.05; Figure 1C), further confirming the enhancement of autophagic activity under prolonged aldosterone stimulation.

**Figure 1.**
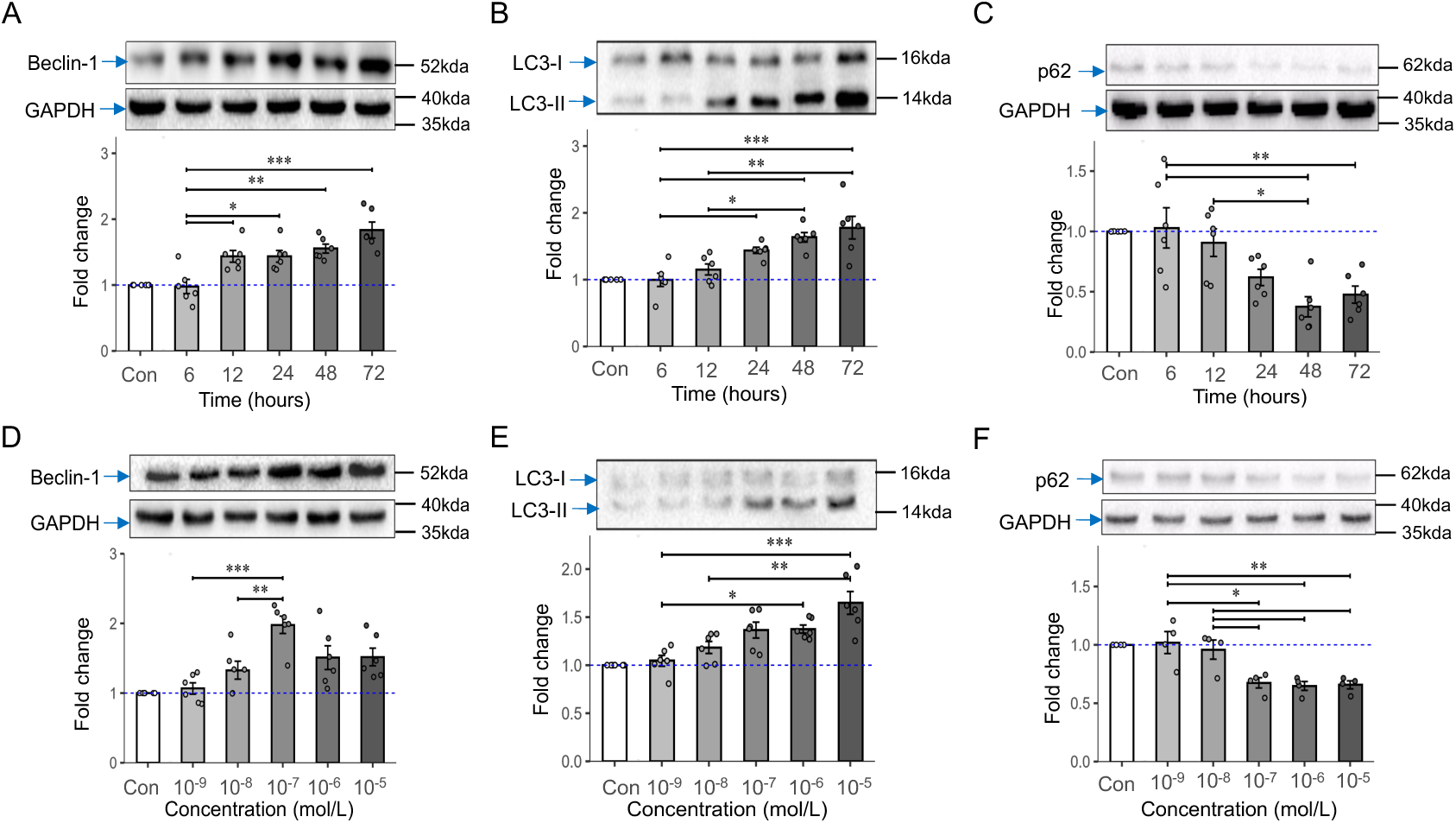
Dose- and time-response of autophagy flux proteins expressions. A) Time-response of Beclin-1 expression. B) Time-response of LC3-I/LC3-II expression. C) Time-response of p62 expression. D) Dose-response of Beclin-1 expression. E) Dose-response of LC3-I/LC3-II expression. F) Dose-response of p62 expression. N=4-7 in each group. *P < 0.05, **P < 0.01, ***P < 0.001.

In the concentration dimension, the expression of autophagic proteins exhibited robust upregulation beginning at 10^-7^ mol/L for Beclin-1 (Ps < 0.05; Figures 1D) and 10^-8^ mol/L (Ps < 0.05; Figures 1E) aldosterone for LC3-II/LC3-I ratio, whereas SQSTM1/p62 levels were already reduced at 10^-7^ mol/L and continued to decline with higher concentrations (Ps < 0.05; Figure 1F). Taken together, these findings delineate a clear dose- (10^-7^ mol/L) and time-dependent (48 h) induction of autophagy in HUVECs in response to aldosterone.

In HUVECs, MR localization was observed in both the cytoplasm and nucleus under basal conditions. Following 48 hours of aldosterone stimulation, MR exhibited a marked translocation from the cytoplasm to the nucleus, resulting in a significant increase in nuclear MR expression (t(12) = –2.390, p = 0.038; Figure 2A, C) and a trend toward decreased cytoplasmic expression (t(12) = – 2.390, p = 0.038; Figure 2B, D). Quantitative immunofluorescence analysis corroborated these findings, revealing enhanced MR signal intensity within the nucleus after aldosterone treatment (t(168) = 0.775, p = 0.456; Figure 2E–F). Lysosomes were predominantly distributed in the perinuclear region under control conditions. Upon aldosterone stimulation, the degree of MR-lysosome co-localization significantly increased (t(92) = –3.728, p < 0.001; Figure 2G–H). Together, these results indicate that excessive aldosterone stimulation drives MR translocation not only to the nucleus-consistent with its canonical transcriptional activity-but also toward lysosomal compartments, suggesting a potential role for lysosome-associated MR in non-genomic or degradative signaling pathways.

**Figure 2.**
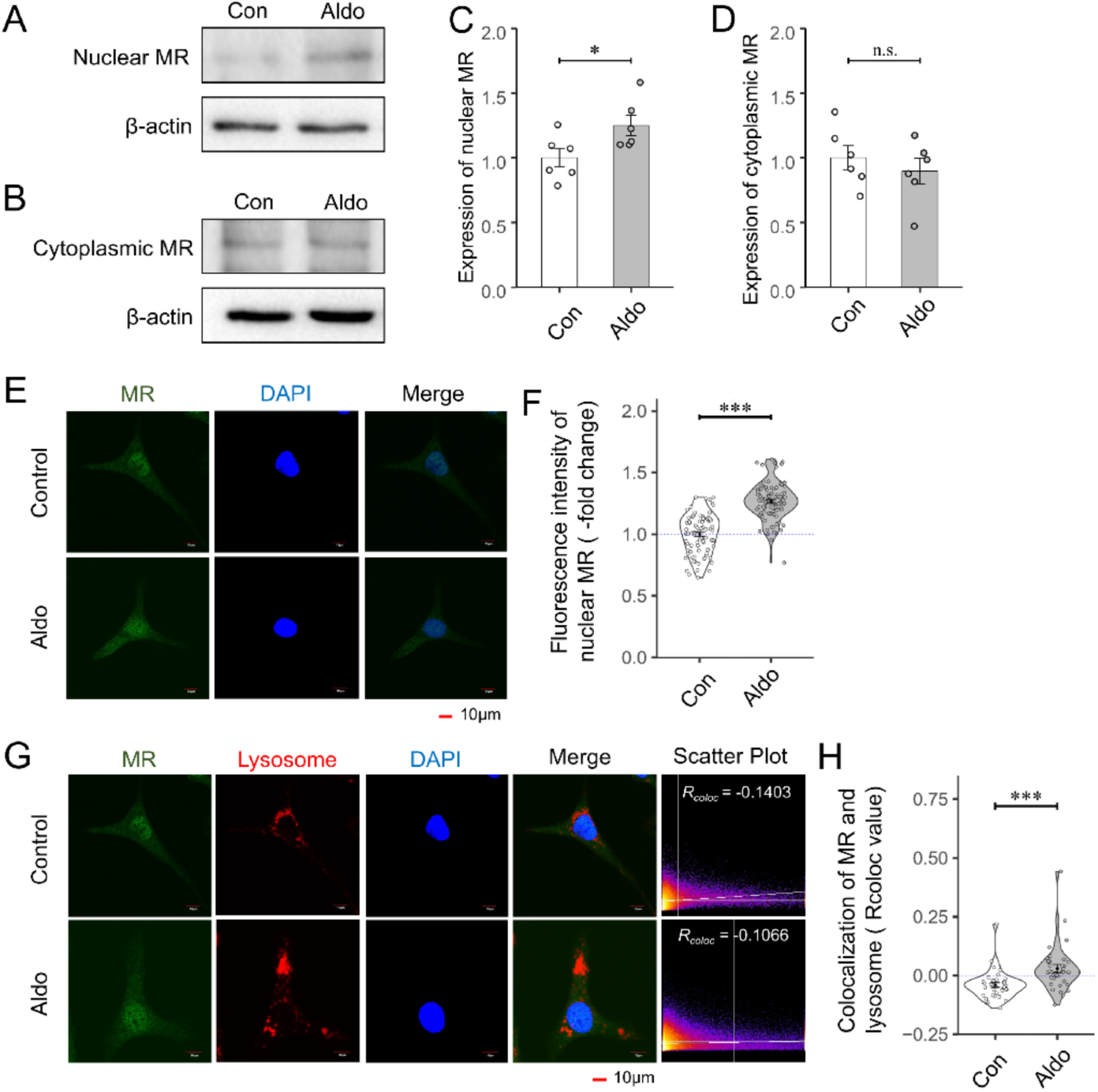
Aldosterone induced MR translocation. A) Representative blot of MR in nucleus. B) Representative blot of MR in cytosol. C) Statistical analysis of MR expression in nucleus (n=6). D) Statistical analysis of MR expression in cytosol (n=6). E) Representative immunofluorescence images of MR expression in nucleus. F) Immunofluorescence intensity analysis of MR expression in nucleus. G) Representative immunofluorescence images of co-localization of MR and lysosomes. H) Rcoloc value analysis of co-localization of MR and lysosomes. *P < 0.05, ***P < 0.001.

These results suggest that aldosterone activates the autophagy pathway in HUVECs. However, eplerenone (Epl, MR antagonist), 3-methyladenine (3-MA, autophagy inhibitor) and tempol (Temp, antioxidant) significantly suppressed the aldosterone-induced upregulation of Beclin-1 (Epl: t(15)*_Aldo>Aldo+Epl_* = 3.812, p = 0.005, Figure 3A; Temp: t(15)*_Aldo>Aldo+Temp_* = 2.605, p = 0.049, Figure 3G; 3-MA: t(9)*_Aldo> Aldo+3-MA_* = 3.460, p = 0.018, Figure 3J) and the LC3-II/LC3-I ratio (Epl: t(15)*_Aldo>Aldo+Epl_* = 4.345, p = 0.002, Figure 3B; Temp: t(12)*_Aldo>Aldo+Temp_* = 4.887, p = 0.001, Figure 3H; 3-MA: t(12)*_Aldo> Aldo+3-MA_* = 3.657, p = 0.009, Figure 3K), while concurrently promoting the expression of p62 (Epl: t(15)*_Aldo>Aldo+Epl_* = -3.784, p = 0.005, Figure 3C; Temp: t(9)*_Aldo>Aldo+Temp_* = -2.900, p = 0.042, Figure 3I; 3-MA: t(9)*_Aldo>Aldo+3-MA_* = -4.223, p = 0.006, Figure 3L), which was suppressed by aldosterone. In contrast, 5-hydroxydecanoate (5-HD, mitochondrial KATP channel inhibitor) failed to reverse the aldosterone-induced increases in Beclin-1 (t(15)*_Aldo>Aldo+5-HD_* = -0.858, p = 0.674, t(15)*_Aldo>5-HD_* = 3.819, p = 0.004, t(15) *_5- HD >Aldo+5-HD_* = -4.676, p < 0.001, Figure 3D) and the LC3-II/LC3-I ratio (t(15)*_Aldo>Aldo+5-HD_* = 0.009, p = 1.000, t(15)*_Aldo>5-HD_* = 3.198, p = 0.016, t(15) *_5- HD >Aldo+5-HD_* = -3.188, p = 0.016, Figure 3E), and was unable to restore p62 to normal expression levels (t(15)*_Aldo>Aldo+5-HD_* = 0.296, p = 0.953, t(15)*_Aldo>5-HD_* = - 2.964, p = 0.025, t(15) *_5-HD >Aldo+5-HD_* = 3.261, p = 0.014, Figure 3F). Furthermore, rapamycin (Rapa, autophagy activator) exacerbated the aldosterone-induced autophagy activation, as evidenced by further elevated levels of Beclin-1 (t(12)*_Aldo>Aldo+Rapa_* = -2.629, p = 0.054, t(12)*_Aldo>Rapa_* = 0.104, p = 0.994, t(12) *_Rapa >Aldo+Rapa_* = -2.733, p = 0.045, Figure 3M) and the LC3-II/LC3-I ratio (t(12)*_Aldo>Aldo+Rapa_* = -3.778, p = 0.007, t(12)*_Aldo>Rapa_* = -2.323, p = 0.091, t(12) *_Rapa >Aldo+Rapa_* = 1.452, p = 0.347, Figure 3N), coupled with a further reduction in p62 expression (t(15)*_Aldo>Aldo+Rapa_* = 4.855, p < 0.001, t(15)*_Aldo>Rapa_* = 4.043, p = 0.003, t(15) *_Rapa >Aldo+Rapa_* = 0.812, p = 0.701, Figure 3O). These results suggest that aldosterone may regulate autophagy pathway via oxidative stress.

**Figure 3.**
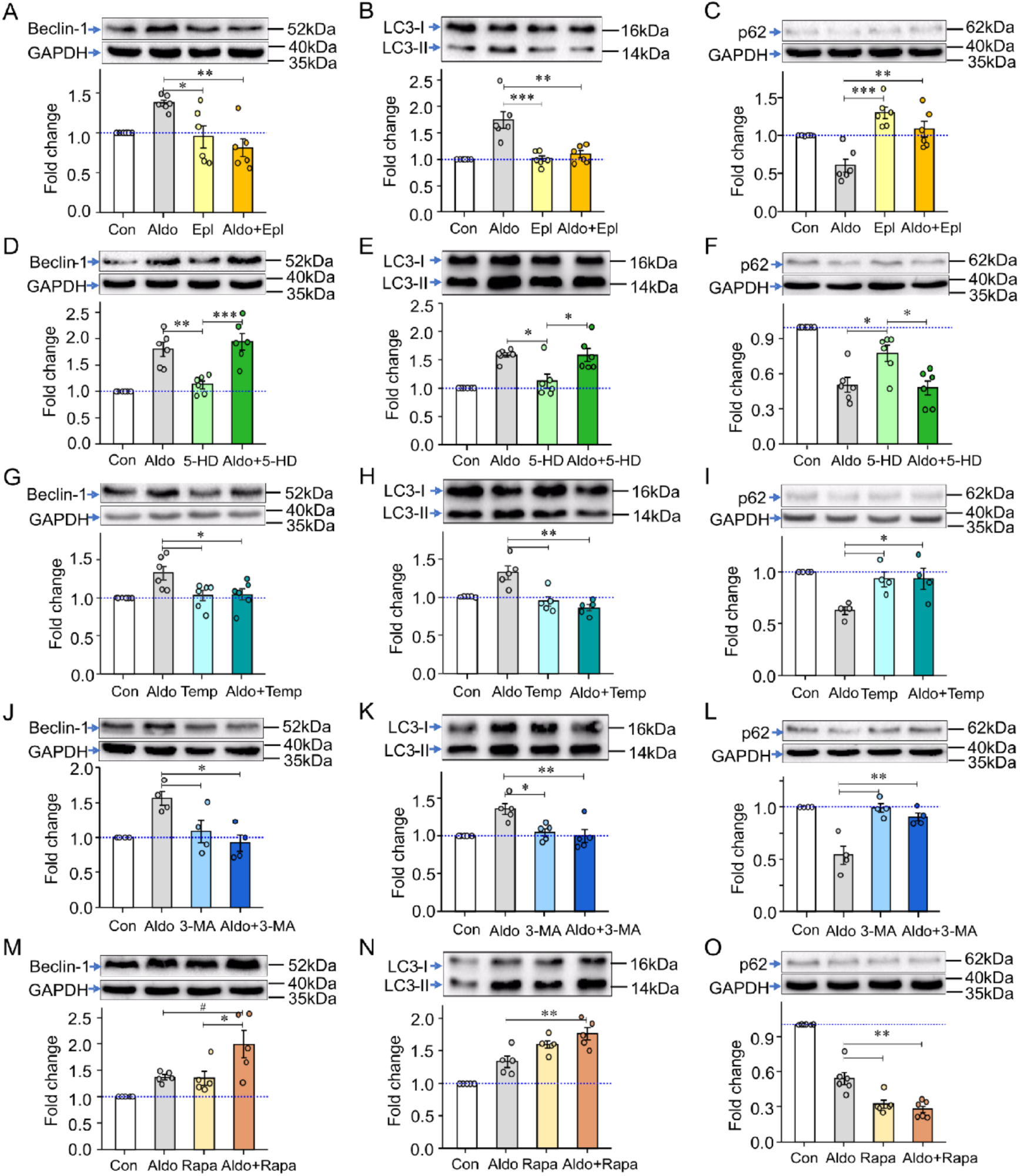
Signal pathways involved in aldosterone induced autophagy flux. The effect of A-C) eplerenone (Epl, 10^-7^ mol/L), D-F) 5-hydroxydecanoate (5-HD, 10^-4^ mol/L), G-I) tempol (Temp, 10^-5^ mol/L), J-L) 3-methyladenine (3-MA, 2×10^-3^mol/L) and M-O) rapamycin (Rapa, 10 mg/L) on aldosterone-induced autophagy flux. N=4-6 in each group. *P < 0.05, **P < 0.01, ***P < 0.001.

Aldosterone stimulation markedly enhanced the interaction between Beclin-1 and the mineralocorticoid receptor. As shown in Figure 4, confocal co-localization analysis revealed a significant increase in Beclin-1-MR co-localization following aldosterone treatment (Figure 4A). Consistently, co-immunoprecipitation assays confirmed that aldosterone exposure strengthened the physical association between Beclin-1 and MR (Figure 4B). These findings suggest that Beclin-1 may facilitate the recruitment of MR into the autophagic machinery, promoting its subsequent trafficking toward lysosomal degradation.

**Figure 4.**
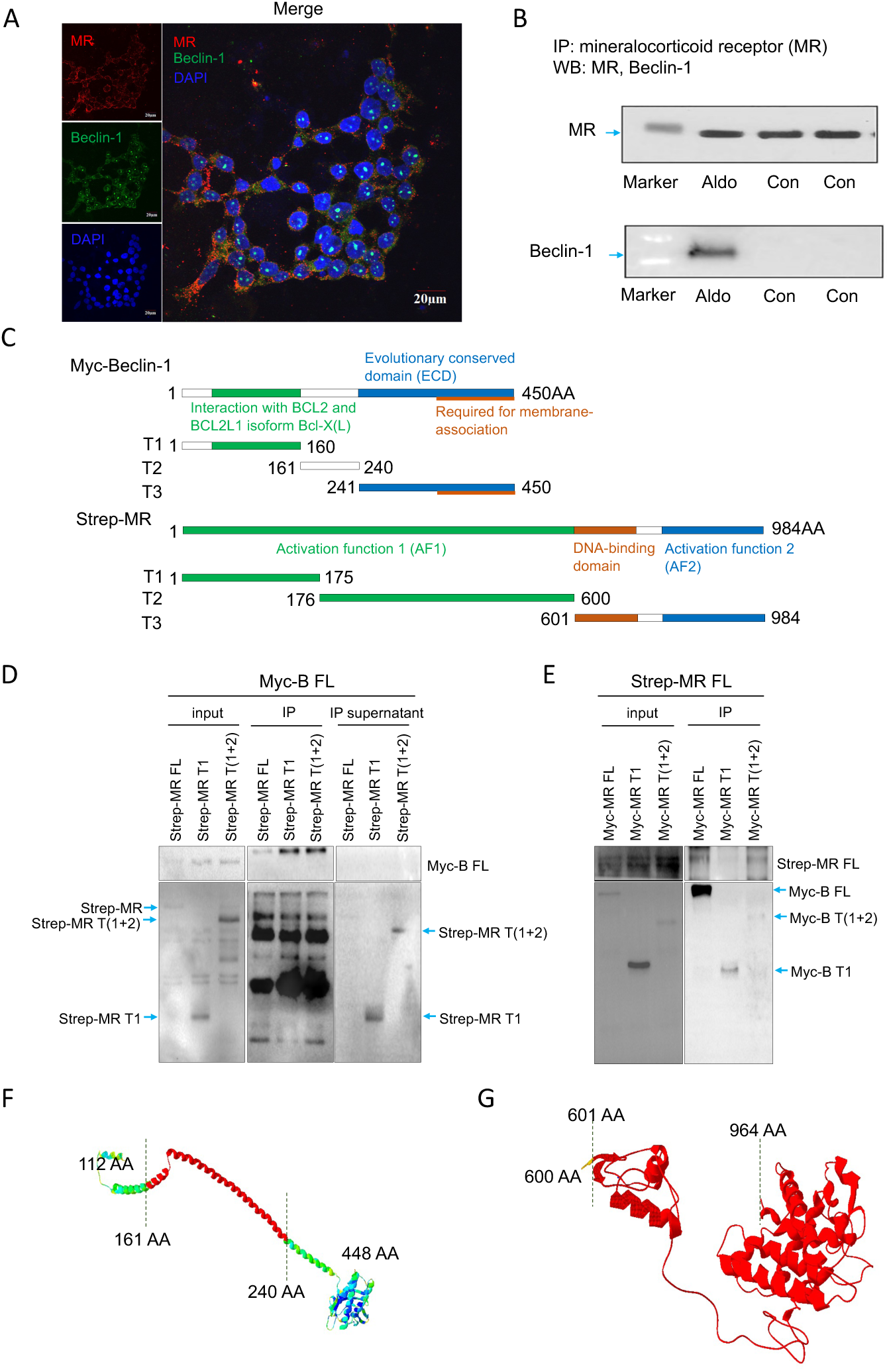
Aldosterone induced interaction of Beclin-1 and MR. A) Representative immunofluorescence image of co-localization of Beclin-1 and MR. B) Representative blot image of co-immunoprecipitation of Beclin-1 and MR. C) Illustration of three truncated parts of Beclin-1 and MR linked with Myc or Strep tags. D) Representative blot image of co-immunoprecipitation of truncated T1, T (1+2) and intact MR with Beclin-1. E) Representative blot image of co-immunoprecipitation of truncated T1, T (1+2) and intact Beclin-1 with MR. F, G) The predicted binding regions between Beclin-1 and MR proteins using SWISS-MODEL software are marked in red. Myc-Beclin-1 T2 (161-241 AA) in **F**, Strep-MR T3 (600-964 AA) in **G**.

To map the specific regions mediating this interaction, three distinct truncation constructs of each protein were generated and expressed in HEK293T cells (Figure 4C). Co-immunoprecipitation analysis demonstrated that the Beclin-1 T2 fragment (amino acids 161-241) directly bound to the MR T3 fragment (amino acids 601-984) (Figure 4D, E). These results pinpoint the interaction interface to the central domain of Beclin-1 and the C-terminal region of MR, providing molecular evidence that Beclin-1 may act as a scaffold linking MR to the autophagic degradation pathway under aldosterone stimulation.

To investigate the structural characteristics of the target proteins, the three-dimensional (3D) structures were predicted using the SWISS-MODEL software, and the obtained modeling results are presented in Figure 4F and Figure 4G. Specifically, Figure 4F displays the predicted 3D structure of the Myc-Beclin-1 T2 fragment. In this figure, the region highlighted in red corresponds to the Myc-Beclin-1 T2 fragment, which spans the amino acid (AA) sequence range of 161-241 AA. This red-highlighted structure provides a clear visualization of the spatial conformation of the Myc-Beclin-1 T2 fragment, laying a foundation for subsequent analyses of its structural features and potential functional sites.

In contrast, Figure 4G focuses on the predicted 3D structure of the Strep-MR T3 fragment. This figure explicitly shows the spatial arrangement of the Strep-MR T3 fragment, which covers the AA sequence from 600-964 AA. The detailed structural information of the Strep-MR T3 fragment presented in Figure 4G is crucial for further exploring its interaction mechanisms with other molecules or its role in related biological processes.

Collectively, these SWISS-MODEL-predicted structures (Figure 4F, G) offer reliable structural references for in-depth studies on the functions and molecular mechanisms of the Myc-Beclin-1 T2 (161-241 AA) and Strep-MR T3 (600-964 AA) fragments.

To explore the downstream inflammatory response driven by MR activation, we next examined potential MR binding sites within the promoters of key proinflammatory cytokine genes. Using JASPAR and NCBI databases, putative MR response elements were identified in the promoter regions of *IL-1β*, *IL-6*, and *TNF-α* (Figure 5A-C). Chromatin immunoprecipitation followed by quantitative PCR (ChIP-qPCR) confirmed that aldosterone stimulation significantly increased MR occupancy at these promoter regions. Specifically, the promoter sequences of *IL-1β* (t(4) = –9.951, p = 0.001), *IL-6* (t(4) = –3.393, p = 0.041), and *TNF-α* (t(4) = –18.185, p < 0.001) were markedly enriched in the anti-MR immunocomplex following aldosterone exposure (Figure 5D-I). These findings demonstrate that aldosterone-activated MR directly engages with the promoters of proinflammatory cytokines, initiating a transcriptional cascade that amplifies the endothelial inflammatory response.

**Figure 5.**
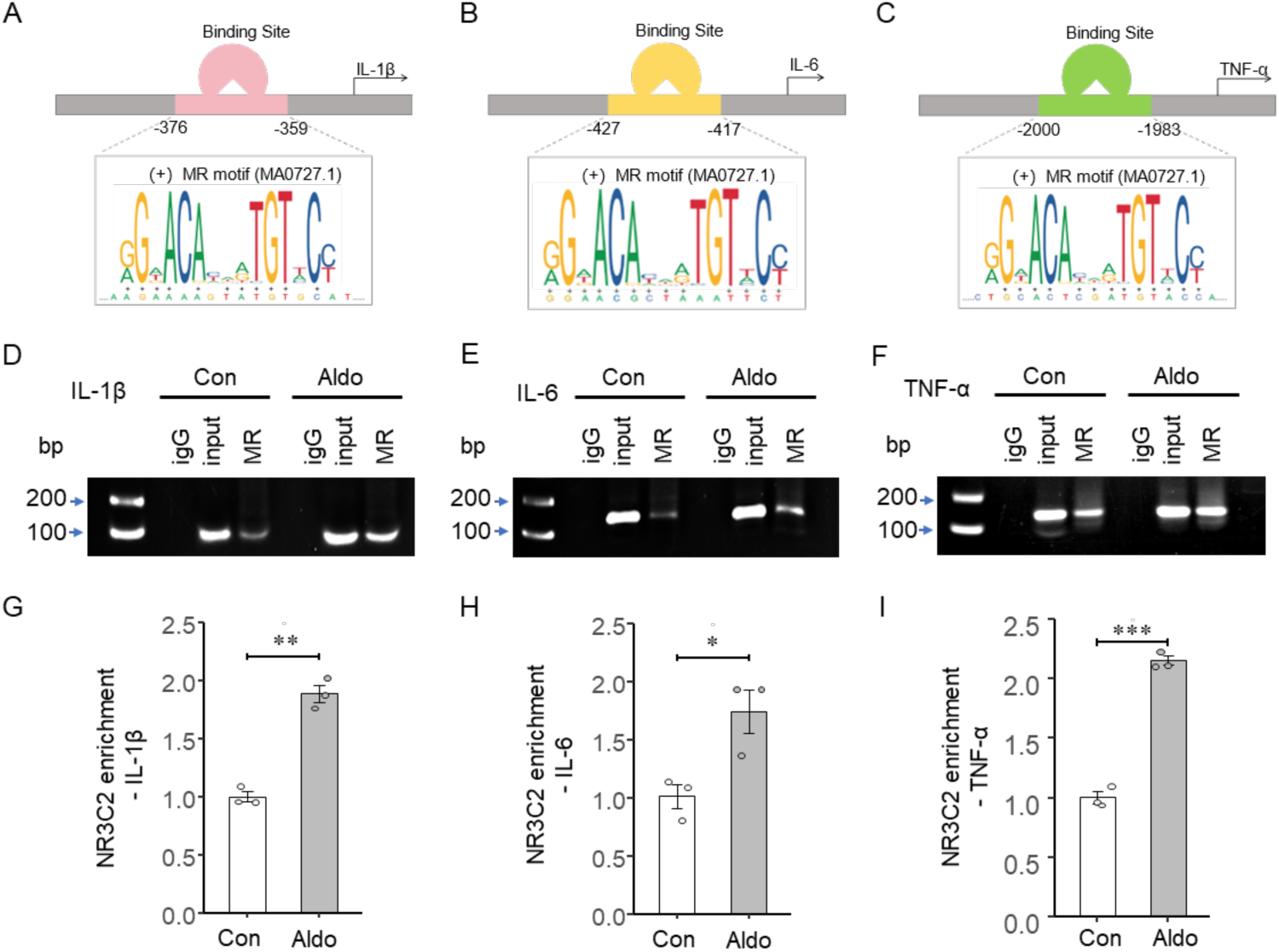
Aldosterone stimulation activates downstream signal. (A-C) Schematic diagrams of the predicted binding sites between the human *IL-1β* promoter, the human *IL-6* promoter, the human *TNF-α* promoter and the MR transcription factor through bioinformatics. (D-F) Agarose gel electrophoresis images of the enrichment of MR with the promoters of *IL1β*, *IL-6*, and *TNF-α* in HUVEC cells determined by ChIP assay. (G-I) Quantitative analysis of *IL1β*, *IL-6*, and *TNF-α* promoter expression by ChIP-qPCR. N=3 in each group. *P < 0.05, **P < 0.01, ***P < 0.001.

Overexpression of Beclin-1 (encoded by the *BECN1* gene) in HUVECs was confirmed by immunoblot analysis (t(4) = 4.479, p = 0.011; Figure 6A). Notably, *BECN1* overexpression markedly attenuated aldosterone-induced MR translocation to the nucleus (t(208) = 17.337, p < 10⁻⁵; Figure 6B), suggesting that elevated *BECN1* levels restrain MR’s nuclear signaling. Consistent with this, ChIP-qPCR analysis revealed that *Beclin-1* overexpression significantly reduced MR enrichment at the promoters of *IL-1β* (t(4) = 12.798, p < 0.001), *IL-6* (t(4) = 7.533, p = 0.002), and *TNF-α* (t(10) = 2.950, p = 0.015) in response to aldosterone stimulation (Figure 6C-H), indicating diminished transcriptional activation of proinflammatory genes.

**Figure 6.**
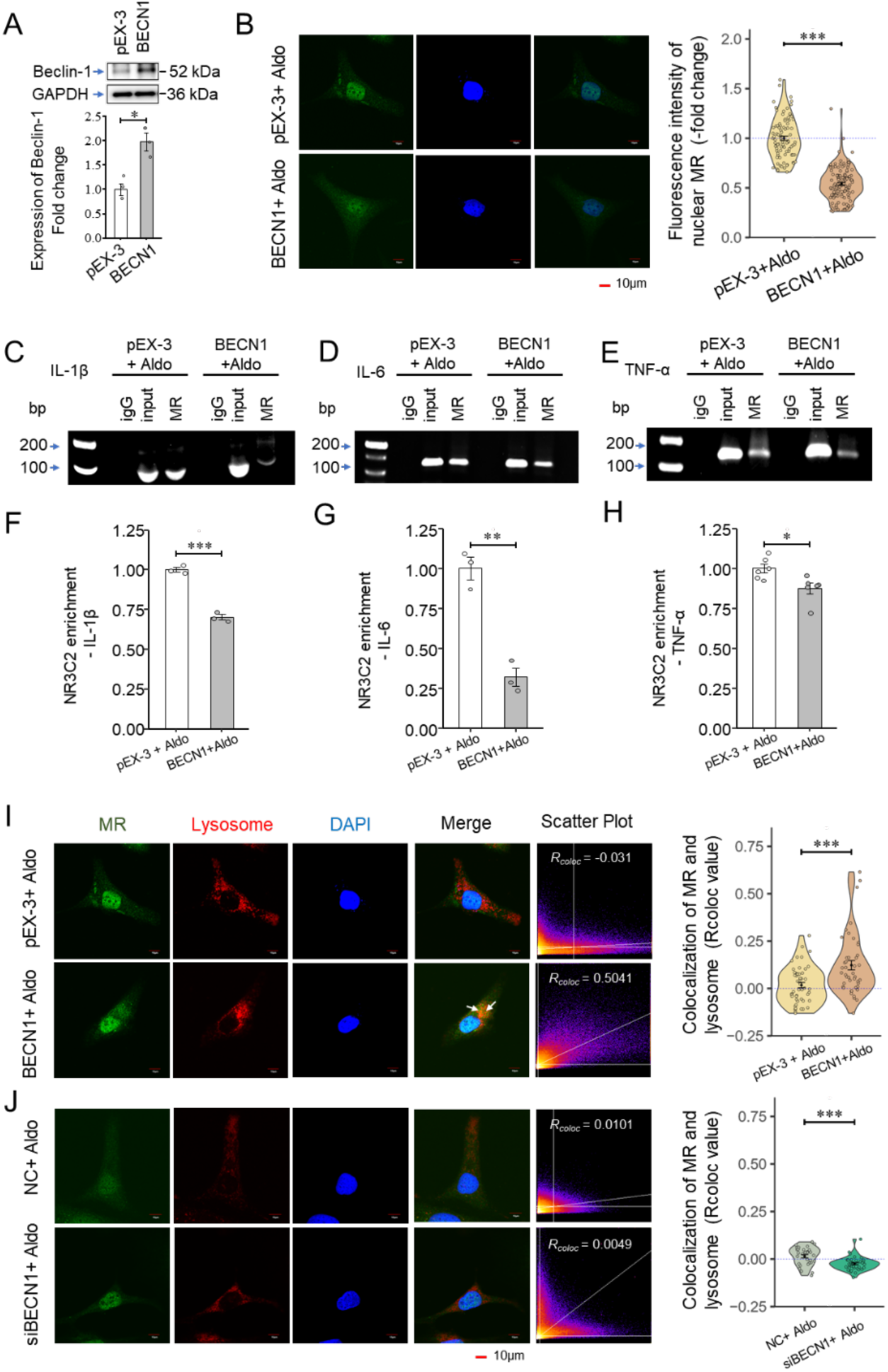
Effect of Beclin-1 manipulation. A) Determination of Beclin-1 over-expression in HUVECs. B) Representative immunofluorescence images of MR expression in nucleus, immunofluorescence intensity analysis of MR expression in nucleus. C-E) Agarose gel electrophoresis images of the enrichment of MR with the promoters of *IL1β, IL-6*, and *TNF-α* in HUVEC cells determined by ChIP assay. (F-H) Quantitative analysis of the enrichment of NR3C2 (MR) with the promoters of IL1β, IL-6, and TNF-α in HUVEC cells determined by ChIP assay in C-E. Representative immunofluorescence images of co-localization of MR and lysosomes; Rcoloc value analysis of co-localization of MR and lysosomes in (I) *BECN1* over-expression and (J) *BECN1* knockdown HUVECs with Aldo treatment. *P < 0.05, **P < 0.01, ***P < 0.001.

At the subcellular level, *BECN1* overexpression enhanced the co-localization of MR with lysosomes (t(92) = –3.728, p < 0.001; Figure 6I), whereas *BECN1* knockdown via siRNA abolished this effect (t(65) = 3.802, p < 0.001; Figure 6J). Together, these findings suggest that elevated Beclin-1 diverts MR from its canonical nuclear route toward lysosomal compartments, thereby promoting MR sequestration and degradation. This mechanism may represent a critical checkpoint through which autophagic activity restrains excessive MR-mediated inflammatory signaling under aldosterone stimulation.

To further investigate the role of *Beclin-1* in vivo, we employed transgenic *Becn1*-overexpressing (*Becn1*-tg) mice subjected to chronic aldosterone stimulation. Remarkably, Becn1-tg mice exhibited significantly lower systolic blood pressure beginning in the second week of aldosterone treatment and persisting thereafter, compared with wild-type (WT) controls (Ps < 0.001), while body weight remained comparable between groups (Ps > 0.05; Figure 7A-D). Histological assessment revealed that aldosterone-induced vascular remodeling was markedly attenuated in *Becn1*-tg mice. Specifically, the aortic medial cross-sectional area was significantly smaller in Becn1-tg mice than in aldosterone-treated WT counterparts (t(8)*_WT+Aldo > Becn1-tg_* = 9.400, p < 0.001; t(8)*_WT > WT+Aldo_* = –11.279, p < 0.001; t(8)*_WT > Becn1-tg+Aldo_* = –3.375, p = 0.018; t(8)*_WT+Aldo > Becn1-tg+Aldo_* = 7.904, p < 0.001; Figure 7E–F). Similarly, indices of cardiac hypertrophy, including ratio of heart weight to body weight (t(16)*_WT+Aldo > Becn1-tg_* = 4.982, p < 0.001; t(16)*_WT > WT+Aldo_* = –6.136, p < 0.001; t(16)*_WT+Aldo > Becn1-tg+Aldo_* = 5.727, p < 0.001; Figure 7G-H), mRNA expressions of BNP (t(32)*_WT > WT+Aldo_* = -9.150, p < 0.001; t(32) *_WT+Aldo > Becn1-tg_* = 11.291, p < 0.001; t(32)*_WT+Aldo > Becn1-tg+Aldo_* = 7.081, p < 0.001; t(32) *_Becn1-tg > Becn1-tg+Aldo_* = -4.210, p = 0.001; Figure 7J) and β-MHC (t(20)*_WT > WT+Aldo_* = -4.324, p = 0.002; t(20) *_WT+Aldo > Becn1-tg_* = 4.868, p < 0.001; t(20)*_WT+Aldo > Becn1-tg+Aldo_* = 3.359, p = 0.015; Figure 7K), were significantly reduced in Becn1-tg mice compared with WT mice following aldosterone exposure.

**Figure 7.**
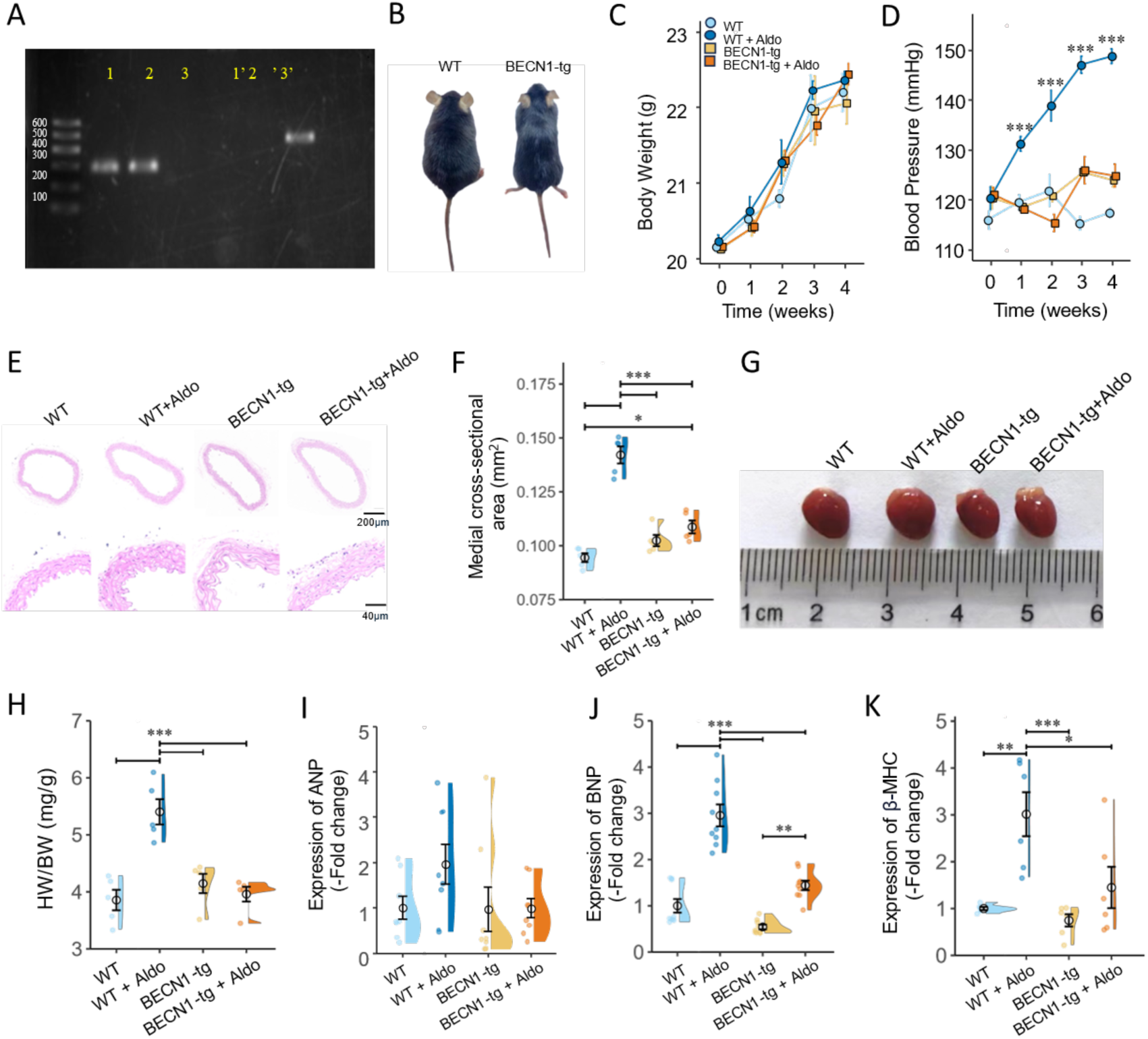
Hypertrophy response to chronic aldosterone stimulation. A) Results of agarose gel electrophoresis identification: 1 and 1’, 2 and 2’ used Becn1+/+ primers, and their target bands were a single band of 283 bp. 3 and 3’ used WT primers, and their target bands were a single band of 453 bp. B) Comparison of phenotype. C) Changes of body weight. D) Changes of systolic blood pressure. E) Representative images of aortic cross-section. F) Statistical analysis of medial cross-section area. G) Representative images of heart. H) Comparison of ratio of heart weight (HW) to body weight (BW). I) mRNA expression of ANP. J) mRNA expression of BNP. K) mRNA expression of β-MHC. *P < 0.05, **P < 0.01, ***P < 0.001.

Together, these in vivo findings demonstrate that Beclin-1 overexpression mitigates aldosterone-induced cardiovascular remodeling, lowering blood pressure and protecting against vascular and cardiac hypertrophy. These results reinforce the concept that Beclin-1-mediated modulation of MR signaling provides a physiological safeguard against mineralocorticoid-driven cardiovascular injury.

To verify the impact of Beclin-1 overexpression on aldosterone stimulation, we firstly tested the MR expression in cytoplasm and nucleus in heart tissue. Results showed that, compared with WT mice, aldosterone treatment significantly reduced cytoplasmic MR levels (t(16)*_WT > WT+Aldo_* = 4.973, p < 0.001, Figure 8A) and increased nuclear MR (t(16)*_WT > WT+Aldo_* = -2.969, p = 0.041, Figure 8B), indicating that aldosterone promotes the translocation of MR from cytoplasm to nucleus. In addition, BECN1 overexpression attenuated aldosterone-induced MR nuclear accumulation (t(16)*_WT+Aldo > Becn1-tg+Aldo_* = - 2.955, p = 0.042, Figure 8B) without affecting cytoplasmic MR abundance (t(16)*_WT+Aldo > Becn1-tg+Aldo_* = -0.755, p = 0.873, Figure 8A), suggesting that Beclin-1 overexpression markedly reduced the MR translocation to nucleus (Figure 8A, B). Inflammatory cytokines IL-6 (t(24) *_WT+Aldo > Becn1-tg_* = 3.239, p = 0.017; t(24)*_WT+Aldo > Becn1-tg+Aldo_* = 3.099, p = 0.024; Figure 8D), TNF-ɑ (t(24)*_WT > WT+Aldo_* = -4.778, p < 0.001; t(24) *_WT+Aldo > Becn1-tg_* = 3.078, p = 0.025; t(24)*_WT+Aldo > Becn1- tg+Aldo_* = 2.967, p = 0.032; Figure 8E) mRNA expressions were significantly lower in TG mice heart compared with that in wild type mice, but not IL-1β (ANOVA: F(3,16) = 3.135, p = 0.055; Figure 8C). Serum levels of IL-1β (t(28)*_WT > WT+Aldo_* = -4.037, p = 0.002; t(28)*_WT+Aldo > Becn1-tg+Aldo_* = 5.8071, p < 0.001; t(28) *_Becn1-tg > Becn1-tg+Aldo_* = 4.152, p = 0.002; Figure 8F), IL-6 (t(16)*_WT > WT+Aldo_* = -4.077, p = 0.004; t(16) *_WT+Aldo > Becn1-tg_* = 4.858, p < 0.001; t(16)*_WT+Aldo > Becn1-tg+Aldo_* = 4.847, p < 0.001; Figure 8G) and TNF-ɑ (t(28)*_WT > WT+Aldo_* = -4.488, p < 0.001; t(28)*_WT+Aldo > Becn1-tg+Aldo_* = 4.465, p < 0.001; Figure 8H) were lower in TG mice than that in wild mice. In addition, oxidative markers such as DHE (t(8)*_WT > WT+Aldo_* = -3.824, p = 0.021; t(8) *_WT+Aldo > Becn1-tg_* = 3.915, p = 0.019; t(8)*_WT+Aldo > Becn1-tg+Aldo_* = 4.246, p = 0.012; Figure 8I, J) and MDA (t(16)*_WT+Aldo > Becn1-tg+Aldo_* = 4.060, p = 0.005; Figure 8 K) were also showed similar patterns in TG mice compared with wild type mice.

**Figure 8.**
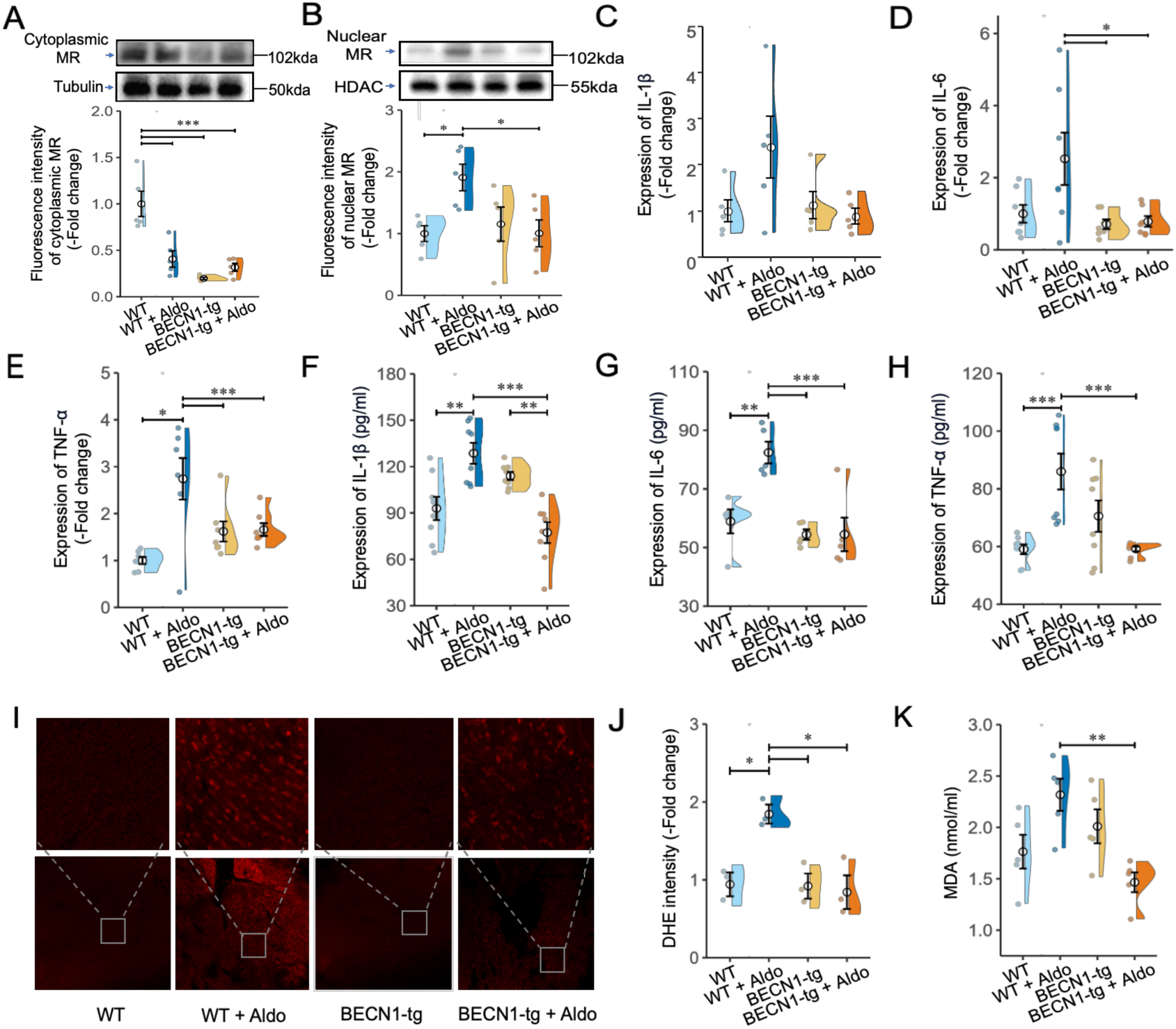
Effects of Beclin-1 over-expression on MR translocation, inflammation, oxidative stress in response to aldosterone stimulation. A) Protein expression of MR in cytoplasm. B) Protein expression of MR in nucleus. C) Heart mRNA expression of *IL-1β*. D) Heart mRNA expression of *IL-6*. E) Heart mRNA expression of *TNF-ɑ*. F) Serum level of IL-1β. G) Serum level of IL-6. H) Serum level of TNF-ɑ. I) Representative heart images of DHE staining. J) Density analysis of DHE. K) serum level of MDA. *P < 0.05, **P < 0.01, ***P < 0.001.

## DISCUSSION

Autophagy serves as an evolutionarily conserved catabolic program that maintains cellular homeostasis, adapts to stress, and modulates signal transduction through selective degradation of effector proteins. In the cardiovascular system, dysregulated autophagy is closely implicated in hypertension, vascular remodeling, inflammation, and cardiac hypertrophy^18^; however, the crosstalk between autophagy and aldosterone–mineralocorticoid receptor (MR) signaling remains incompletely understood. Here, we demonstrate that Beclin-1 functions as a pivotal endogenous negative regulator that restrains aldosterone-induced MR signaling via autophagic lysosomal degradation, thereby suppressing nuclear translocation, proinflammatory transcription, and pathological cardiovascular remodeling. Our findings uncover a previously unrecognized autophagy-dependent negative feedback loop that counteracts excessive aldosterone action and confers protection against cardiovascular injury.

### Autophagy is activated as a compensatory adaptive response to aldosterone stimulation

The present study shows that aldosterone triggers a time- and dose-dependent increase in Beclin-1 expression, elevated LC3-II/LC3-I ratio, and reduced SQSTM1/p62 accumulation in HUVECs, indicating enhanced autophagic flux. Pharmacological interventions revealed that this induction was abolished by the MR antagonist eplerenone, the antioxidant tempol, and the autophagy inhibitor 3-MA, but not by the mitochondrial KATP channel inhibitor 5-HD. Moreover, the autophagy activator rapamycin further augmented aldosterone-mediated autophagy activation. These results indicate that aldosterone-induced autophagy depends on MR signaling and oxidative stress, rather than mitochondrial KATP activity, and represents a cell-protective compensatory response.

Previous studies have yielded conflicting results regarding aldosterone and autophagy, with reports showing either stimulatory or inhibitory effects in a cell-and context-dependent manner^19, 20^. Our data reconcile these discrepancies by demonstrating that, in endothelial cells, prolonged aldosterone exposure elicits a pro-survival autophagic program governed by Beclin-1. This adaptive mechanism likely limits cellular damage during sustained hormonal stress. When reinforced by Beclin-1 overexpression, endothelial cells become more resistant to aldosterone-induced inflammation and dysfunction, supporting the notion that autophagy acts as a defensive barrier against MR-driven pathological signaling.

### Beclin-1 directly interacts with MR and targets it for lysosomal degradation

A key mechanistic discovery of this work is the physical interaction between Beclin-1 and MR. Co-immunoprecipitation and confocal imaging confirmed that aldosterone enhances the association between endogenous Beclin-1 and MR. Domain-mapping further revealed that the middle domain of Beclin-1 (amino acids 161–241) binds directly to the C-terminal region of MR (amino acids 601–984). This interaction redirects MR from its canonical nuclear transcriptional pathway toward lysosomal sequestration and degradation.

Beclin-1 is best characterized as a core component of the class III PI3K complex that initiates autophagosome formation. Emerging evidence has expanded its function to that of a scaffold protein regulating the stability and subcellular localization of multiple signaling receptors^21, 22^. Our findings extend this paradigm to nuclear receptors: Beclin-1 binds MR and promotes its lysosomal delivery, thereby reducing the pool of transcriptionally competent receptor. To our knowledge, this is the first report demonstrating that a core autophagy protein directly interacts with MR to limit its signaling output. This mechanism represents a novel post-translational regulatory mode for MR that operates independently of classical transcriptional feedback or kinase-mediated phosphorylation cascades.

### Beclin-1 suppresses aldosterone-induced vascular inflammation by inhibiting MR-driven cytokine transcription

Aldosterone-activated MR is a well-established driver of vascular inflammation that contributes to endothelial dysfunction, hypertension, and target organ damage^23^. Our bioinformatic prediction and ChIP-qPCR analyses identified functional MR-binding elements in the promoters of *IL-1β*, *IL-6*, and *TNF-α*, confirming these proinflammatory genes as direct transcriptional targets of MR. Overexpression of *Beclin-1* markedly reduced MR recruitment to these promoters and suppressed cytokine expression, whereas *Beclin-1* knockdown enhanced MR nuclear accumulation and inflammatory gene expression.

These results indicate that Beclin-1-mediated MR degradation directly blunts the proinflammatory transcriptional program activated by aldosterone. In endothelial cells, excessive inflammation promotes leukocyte adhesion, vascular permeability, and reactive oxygen species production, all of which exacerbate hypertension and remodeling^24^. By targeting MR for lysosomal degradation, Beclin-1 acts as a molecular brake that prevents uncontrolled inflammatory signaling during hyperaldosteronism. This anti-inflammatory mechanism is distinct from canonical MR antagonism and may offer a complementary strategy to reduce vascular inflammation without complete blockade of MR-dependent physiological functions.

### Beclin-1 overexpression in vivo protects against aldosterone-induced hypertension, vascular remodeling, and cardiac hypertrophy

To validate the pathophysiological relevance of our in vitro findings, we employed *Beclin-1* transgenic (*Becn*1-tg) mice subjected to chronic aldosterone infusion. Compared with wild-type littermates, *Becn1*-tg mice exhibited significantly lower systolic blood pressure, reduced aortic medial thickening, and diminished cardiac hypertrophy, as reflected by heart weight/body weight ratio and expression of hypertrophic markers ANP, BNP, and β-MHC. In addition, *Beclin-1* overexpression reduced myocardial and systemic levels of IL-1β, IL-6, and TNF-α, as well as oxidative stress markers DHE and MDA. Body weight was comparable between groups, excluding systemic metabolic alterations as a confounding factor.

These in vivo data confirm that Beclin-1 is a potent protective factor against aldosterone-driven cardiovascular disease. The results support a model in which enhanced Beclin-1-dependent autophagy promotes MR degradation, reduces nuclear signaling, limits inflammation and oxidative stress, and ultimately ameliorates pathological remodeling. Importantly, these findings suggest that boosting Beclin-1 activity selectively targets pathological MR signaling while preserving essential physiological functions in electrolyte and fluid homeostasis.

### Translational implications and therapeutic potential

Primary aldosteronism is one of the most prevalent causes of secondary hypertension and a major risk factor for resistant hypertension, atrial fibrillation, heart failure, and chronic kidney disease^12^. Current standard therapy relies on MR antagonists such as spironolactone and eplerenone, which effectively reduce blood pressure and cardiovascular risk but are limited by side effects including hyperkalemia, gynecomastia, and sexual dysfunction^17, 25^. Incomplete responses in some patients highlight the need for alternative therapeutic targets.

Our findings identify the Beclin-1–MR axis as a novel candidate for treating hyperaldosteronism-related cardiovascular disorders. Strategies that enhance Beclin-1 expression or stabilize the Beclin-1–MR interaction could promote MR degradation and suppress pathological signaling without fully blocking MR activity. Such approaches may retain the beneficial effects of MR in renal electrolyte handling while selectively inhibiting inflammation, oxidative stress, and remodeling. Potential translational strategies include gene therapy, small molecules that enhance Beclin-1-dependent autophagy, or peptide mimetics that mimic the MR-binding domain of Beclin-1. Future studies are warranted to evaluate the safety and efficacy of such interventions in preclinical models and eventually in humans.

### Limitations and future perspectives

Several limitations of this study should be noted. First, the detailed molecular mechanisms governing Beclin-1–MR complex formation—including post-translational modifications such as phosphorylation, ubiquitination, or acetylation that may regulate this interaction remain to be fully characterized. Second, this study focused primarily on endothelial cells; whether Beclin-1 regulates MR signaling in cardiomyocytes, renal tubular cells, vascular smooth muscle cells, or immune cells requires further investigation. Third, the high-resolution structural basis of the Beclin-1–MR interaction remains to be determined; crystallographic or cryo-electron microscopy studies will help define the precise binding interface. Fourth, although we demonstrate that Beclin-1 promotes MR lysosomal localization, the detailed trafficking steps and potential involvement of other autophagy adaptors or cargo receptors require clarification.

Future studies should explore whether the Beclin-1–MR regulatory axis is dysregulated in human primary aldosteronism or hypertensive heart disease. It will also be critical to test whether Beclin-1 enhancers or MR-targeting autophagy adaptors can improve cardiovascular outcomes in preclinical models of hypertension and heart failure. Finally, investigation into whether other nuclear receptors are regulated by similar autophagy-dependent degradation mechanisms may uncover broad principles governing signal integration between autophagy and transcriptional regulation.

In summary, we demonstrate that aldosterone induces autophagy and triggers a direct interaction between Beclin-1 and MR. Beclin-1 targets MR for lysosomal degradation, thereby inhibiting nuclear translocation, proinflammatory gene expression, and pathological cardiovascular remodeling. Beclin-1 overexpression in mice protects against aldosterone-induced hypertension, vascular dysfunction, and cardiac hypertrophy. These findings define a novel autophagy-mediated negative feedback loop that restrains excessive aldosterone signaling and identify Beclin-1 as a potential therapeutic target for cardiovascular diseases associated with hyperaldosteronism (see Figure 9).

**Figure 9.**
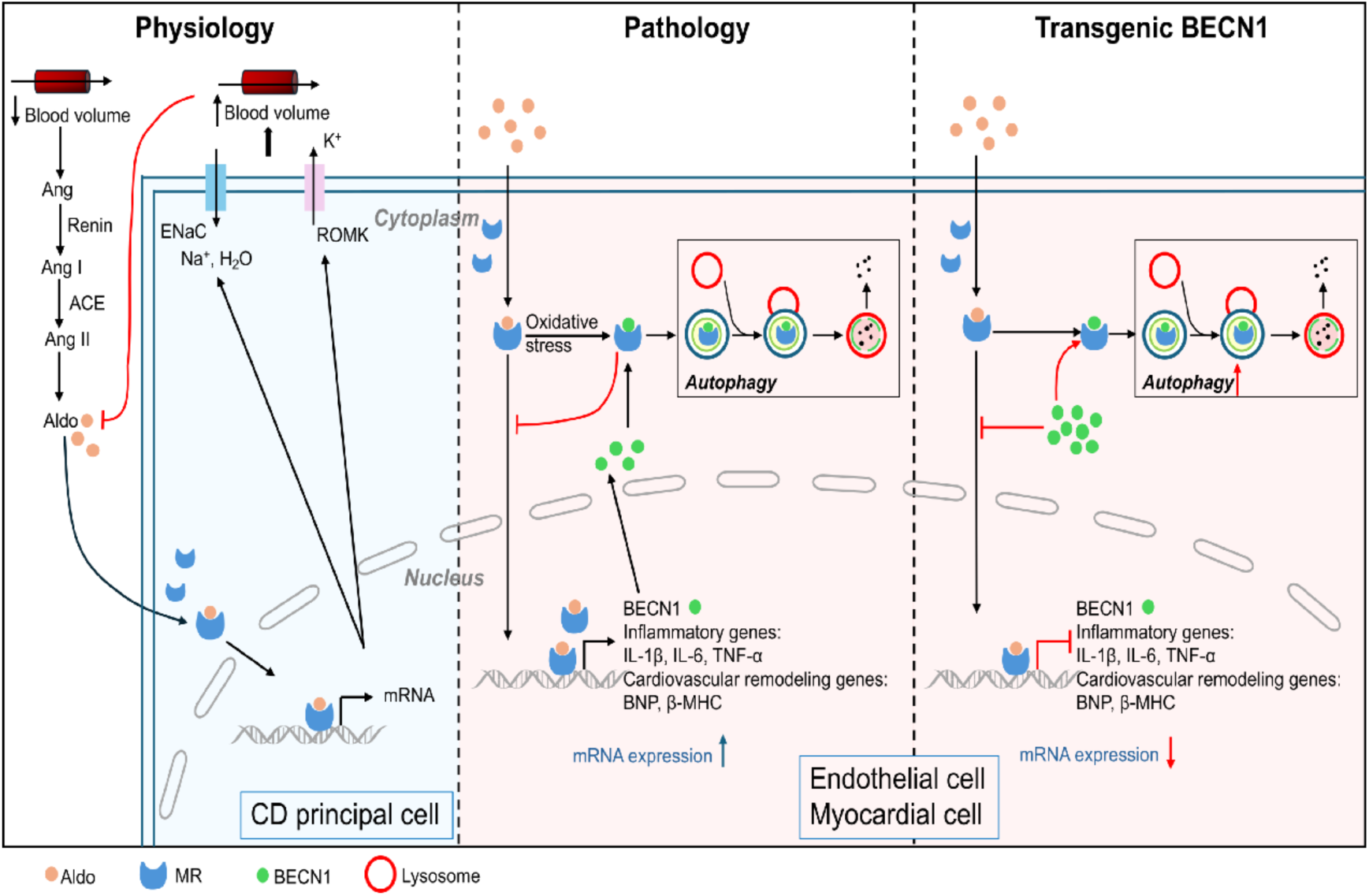
The proposed autophagy-mediated negative feedback loop. Under physiological condition, the renin-angiotensin-aldosterone system (RAAS) is activated. AngiotensinⅡ (AngⅡ) is generated through the action of angiotensin-converting enzyme (ACE) and stimulates the secretion of aldosterone. Aldosterone binds cytoplasmic mineralocorticoid receptor (MR) and induces its nuclear translocation, promoting transcription of ion channels such as ENaC and ROMK. This promotes sodium retention, potassium excretion, and water reabsorption, leading to increased blood volume and trigger negative feedback inhibition of aldosterone production. However, under pathological conditions, aldosterone activates MR signaling in vascular endothelial cells, promoting MR nuclear translocation and transcriptional activation of pro-inflammatory genes, including IL-1β, IL-6, and TNF-α, which contribute to vascular inflammation and cardiovascular remodeling. Concurrently, aldosterone induces autophagy and upregulates the core autophagy regulator Beclin-1. Becline-1 directly interacts with MR, promoting MR sequestration in lysosome-associated compartments and limiting its nuclear accumulation. This Beclin-1-dependent autophagic process establishes an intrinsic negative feedback mechanism that restrains MR signaling and protect against aldosterone-induced cardiovascular injury.

## Acknowledgments

This research was supported by the Start Funding from Nantong First People’s Hospital, Affiliated Hospital of Southeast University (YJRCJJ007), and the National Natural Science Foundation of China (81970422).

## Ethics approval

This study was approved by the Animal Care and Welfare Committee at Soochow University (permission code: 202306A0493), and all animal care and surgical manipulation were guided according to the National Institutes of Health Guidelines for the Use of Laboratory Animals (NIH, Publication number 85-23, revised 1996).

## Author contributions

**Conceptualization:** Guo-Xing Zhang, Koji Murao; **Methodology:** Wen-Yi Jiang, Hao-Tian Zhang, Xiao-Wei Sun, Ya-Mei Gao; **Investigation:** Wen-Yi Jiang, Hao-Tian Zhang, Xiao-Wei Sun, Ya-Mei Gao; **Formal analysis:** Lei Wang, Wen-Yi Jiang, Hao-Tian Zhang, Xiao-Wei Sun, Ya-Mei Gao; **Visualization:** Lei Wang, Wen-Yi Jiang, Hao-Tian Zhang, Xiao-Wei Sun, Ya-Mei Gao; **Writing – original draft:** Lei Wang, Wen-Yi Jiang, Hao-Tian Zhang, Xiao-Wei Sun, Ya-Mei Gao; **Writing – review & editing:** Guo-Xing Zhang, Lei Wang, Koji Murao.

## Data availability

Data will be available on request.

